# A genome-wide RNAi screen identifies host cell cycle regulation as a determinant of *Orientia tsutsugamushi* infection

**DOI:** 10.64898/2026.05.08.723767

**Authors:** Porncheera Chusorn, Yanin Pittayasathornthun, Potjanee Kanchanapiboon, Kittirat Saharat, Kriengkrai Phongkitkarun, Somponnat Sampattavanich, Jeanne Salje

**Author notes:** These authors contributed equally (Porncheera Chusorn, Yanin Pittayasathornthun, Potjanee Kanchanapiboon).

## Abstract

*Orientia tsutsugamushi* (Ot) is an obligate intracellular bacterium that causes scrub typhus, a potentially life-threatening disease. To systematically identify host factors regulating early stages of infection, we performed a microscopy-based genome-wide siRNA screen in HeLa cells. This approach identified 2,989 genes grouped into 55 functional networks that modulate bacterial entry and intracellular translocation. In addition to confirming previously described pathways, including endocytosis and microtubule-dependent trafficking, the screen revealed an association between Ot infection and host cell cycle regulation. We found that Ot preferentially infects and/or replicates in host cells in the S and G2 phases, where intracellular bacterial accumulation is increased relative to G1. Early infection was associated with a shift in host cell cycle distribution, consistent with a delay in progression through S and G2 phases. Longitudinal analysis further showed that these cell cycle states support enhanced bacterial expansion. In parallel, infected cells exhibited reduced proliferation compared to uninfected cells, suggesting that Ot infection alters host cell division dynamics. Together, these findings support a model in which host cell cycle state influences susceptibility to Ot infection and intracellular growth. This work provides a systems-level map of host pathways involved in early infection and identifies cell cycle regulation as an important component of host–pathogen interactions in scrub typhus.

**Author Summary:** Scrub typhus is a potentially life-threatening disease caused by the bacterium *Orientia tsutsugamushi*, which can only survive and replicate inside human cells. Although some host factors involved in infection have been identified, many remain unknown. In this study, we used a large-scale screening approach to systematically identify human genes that influence the bacterium’s ability to enter and move within host cells. Our analysis uncovered multiple pathways required for infection, including a role for the host cell cycle. We found that *O. tsutsugamushi* preferentially accumulates in cells during specific stages of the cell cycle, particularly when cells are preparing to divide. At the same time, infection slows host cell division, suggesting that the bacterium alters the cellular environment to support its own growth. These findings provide new insight into how *O. tsutsugamushi* interacts with human cells and identify potential host processes that could be targeted to limit infection.

## Introduction

Ot is an obligate intracellular Gram-negative bacterium in the order Rickettsiales^1,2^. It causes the severe vector-borne human disease scrub typhus, which is transmitted by *Leptotrombidium* mites^1^. Scrub typhus is historically known to be endemic in Asia, a region encompassing two-thirds of the world’s population. However, recent discoveries of new species of *Orientia* from Latin America^3,4^ and the Middle East^5^ suggest a more global distribution^1^. Infection begins with inoculation of bacteria into the skin at the site of mite feeding, which can develop into a necrotic lesion called an eschar. Bacteria disseminate through the body via blood and lymphatic circulatory systems, where they can infect multiple organs, including the lung, liver, spleen, kidney, heart, and brain. In severe scrub typhus, this can lead to multiple organ failure and death. The exact cell tropism at different stages of human infection is difficult to ascertain. However, eschar biopsies revealed Ot localized in dendritic cells and macrophages^6^, whilst human autopsy data demonstrated Ot infection of endothelial cells^7^. Ot can infect a range of phagocytic and non-phagocytic cells *in vitro,* including fibroblasts, endothelial cells, epithelial cells, monocytes, and macrophages. Since Ot also infects mite salivary and ovary cells, this indicates that, unlike some human-adapted pathogens, the interactions between Ot and host cells are likely based mainly on widely conserved receptors and gene pathways in the host cells.

The early stages of Ot infection have been described through numerous studies. Ot attaches to host cells via bacterial surface proteins, including TSA56^8,9^, ScaB^10^, and ScaC^11^, and interacts with host cell surface proteins, including the proteoglycan Syndecan-4^12^ and fibronectin^8^. Ot then interacts with integrin α5ß1^9^ and induces its entry using clathrin-mediated endocytosis^13^. Studies leading to this model were carried out using intracellular Ot harvested from infected cells. By contrast, Ot that has budded off the surface of infected host cells enters subsequent cells using macropinocytosis rather than clathrin-mediated endocytosis and entry of this bacterial population does not require the involvement of bacterial surface proteins TSA56 or ScaC^14^. Following entry into host cells, Ot escapes from late endosomes^13^, whereupon it induces and evades autophagy through unknown mechanisms^15,16^. Ot lacks flagella and traffics through cells using dynein-dependent motility along microtubules^17,18^. Ot moves to the perinuclear region, where it replicates in a tight microcolony^19^. Whilst its attachment and entry are completed within 30 minutes of infection, and endosomal escape within about 2 hours^13^, bacterial replication begins about 48 hours post infection^20^. We sought to identify host machinery involved in these early stages of infection, prior to the initiation of bacterial replication, using a genome-wide RNAi approach.

The relationship between Ot and the host cell cycle is not well characterised although it was recently shown that Ot strain Ikeda modulates p53 and inhibits cell cycle progression past S phase^21^. The facultative intracellular bacterium *Salmonella enterica* serovar Typhimurium preferentially infects HeLa and RPE-1 cells during the mitotic stages of cell division^22^. It was shown that this is due to increased cholesterol levels in the cell membrane during mitosis, which facilitates bacterial entry^22^. Some bacteria and viruses have been shown to manipulate progression through the host cell cycle using secreted nucleomodulins^23,24^.

RNAi screens involve knocking down individual host cell genes to determine their effect on a particular readout such as infection^25–28^. We designed a microscopy-based screen to quantify the effect of gene knockdown on overall Ot infection levels and the positions and numbers of individual bacteria. We carried out a genome-wide RNAi screen of Ot clinical isolate strain UT76^29^ infection in human epithelial HeLa cells. Our screen identified 55 functional gene networks whose perturbation either enhanced or inhibited Ot entry and/or translocation. In addition to confirming previously implicated pathways, this unbiased analysis uncovered a link between Ot infection and host cell cycle regulation, suggesting that host cell state may play a critical role in determining susceptibility to infection.

## Results

### A Genome-Wide RNAi Screen Identifies Host Factors Regulating Ot Infection

We designed a genome-wide RNAi phenotypic screen to identify host cell genes involved in the early stages of Ot infection (Fig. 1A). Using the clinical Ot strain UT76, isolated from a patient in Thailand^29^, and the human cervical epithelial cell line HeLa, selected for its susceptibility to Ot infection and ease of transfection, we reverse-transfected cells with pools of three siRNAs targeting 20,655 human genes in triplicate. Each plate included positive controls (Kif11, which causes cell death when depleted) and negative controls (scrambled siRNA), as illustrated in the plate map (Supp. Fig. 1A). After 48 hours of siRNA transfection, HeLa cells were infected with Ot at a nominal MOI of 200:1 for an additional 30 hours, resulting in an average of 2.3 bacteria per infected cell. This discrepancy between the number of bacteria added and the number detected in infected cells is typical in Ot infection. This reflects the low percentage of viable bacteria during infection in the lab. The cells were subsequently fixed and stained: Evans blue labeled the cytosol, Hoechst stained the nuclei, and antibodies against the major Ot surface protein TSA56 were used to detect the bacteria (Fig. 1B).

**Figure 1:**
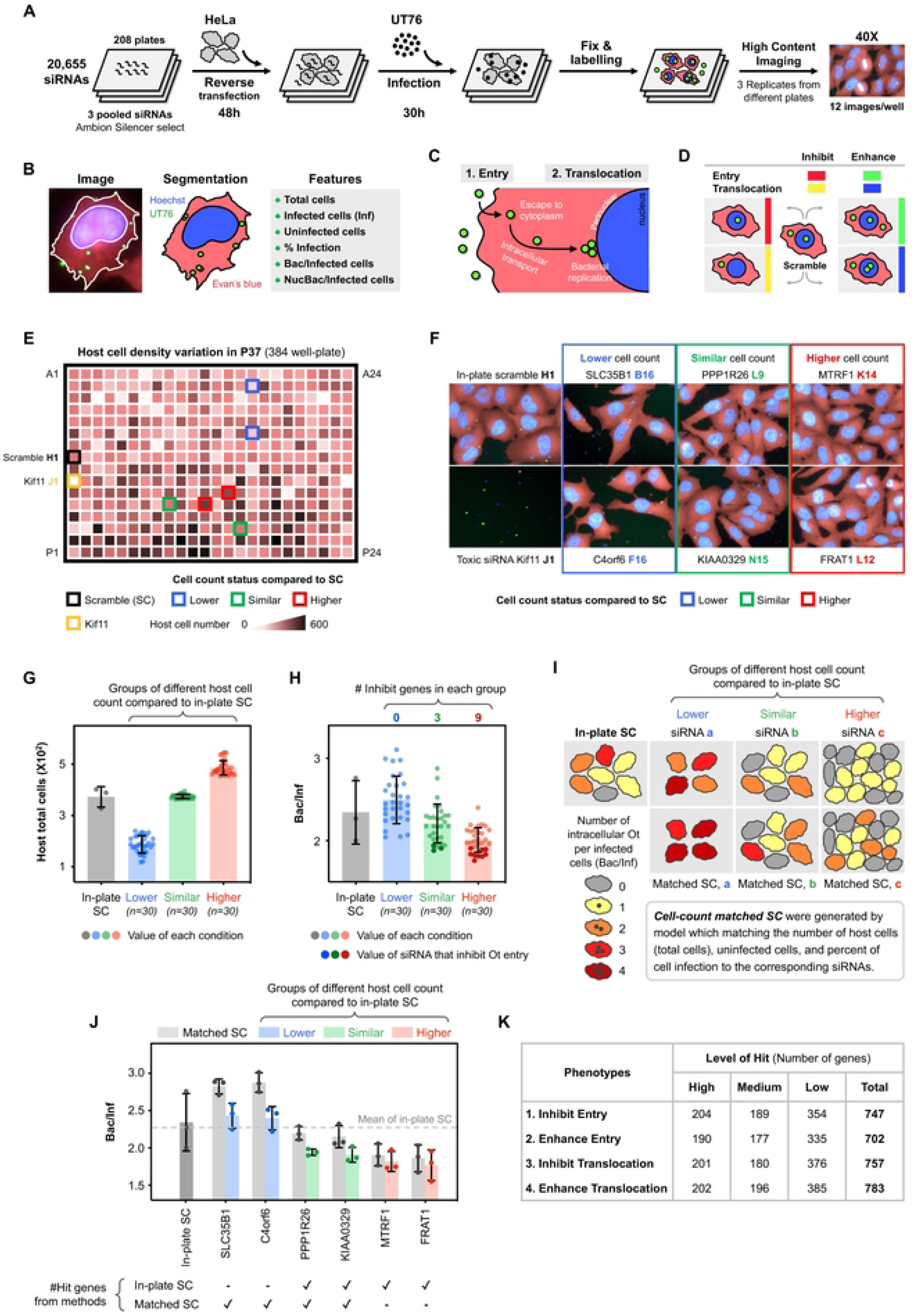
Genome-wide RNAi screen to identify host cell factors involved in Ot entry and translocation. **A.** Schematic overview of screen design. **B.** Representative image showing segmentation of Ot, the nucleus, and the cell outline. Ot is labeled with a rat anti-TSA56 antibody (green), the HeLa cell nucleus is stained with Hoechst (blue), and the cell outline is labeled with Evans blue (red). Features used in our analysis are indicated. % Infection denotes the percentage of host cells that are infected (infected cells/total host cells × 100). Bac/Inf represents the number of bacteria per infected cell, and NucBac/Inf indicates the number of bacteria located in the perinuclear region of the host cell per infected cell. **C.** Overview of the early stages of Ot infection. Two phenotypes were analyzed: “Entry,” which measures the number of bacteria entering host cells (Bac/Inf), and “Translocation,” which assesses the ability of bacteria to traffic to the perinuclear region (NucBac/Inf). **D.** Overview of hit phenotypes and the corresponding color codes used in later figures. Hits that inhibit entry are shown in red, hits that enhance entry in green, hits that inhibit translocation in yellow, and hits that enhance translocation in blue. A schematic illustrating these four phenotypes is provided. **E.** Illustration of host cell number variation at 78 hours post-screening, depicted on a 384-well plate map for siRNA plate P37. The map compares cell count densities between individual siRNAs and the in-plate scramble control. Kif11, a positive control for cell death, is included. **F.** Images from siRNA plate P37 corresponding to panel 1E, showing the 384-well format. Host cell densities were classified into three groups based on their final densities relative to the in-plate scramble: lower (blue), similar (green), and higher (red). **G.** Distribution of total host cell counts in siRNA conditions (n = 30 per group) compared to the in-plate scramble from plate P37. **H.** Distribution of Bac/Inf per condition, stratified by the host cell density groups from panel 1G. The darker color shades indicate siRNAs that significantly reduced Ot entry. Notably, higher host cell density correlated with fewer bacteria per infected cell, indicating that Ot infectivity depends on host cell density. **I.** Development of cell-count-matched scramble controls by adjusting parameters such as total host cell number and percentage of infected cells. These matched scrambles were generated for each siRNA to account for variability across three distinct cell density categories. **J.** Distribution of Bac/Inf in siRNA conditions compared with both in-plate scrambles and cell-count-matched scrambles from plate P37. The use of matched scrambles allowed the identification of additional siRNAs with significant changes in Ot infectivity. **K.** Summary of the number of hits for the four phenotypes across three threshold levels (High, Medium, and Low), which were subsequently used for pathway analysis in Figures 2 and 3.

Automated image analysis was used to delineate host cell boundaries, nuclei, and Ot. From each experimental condition, we extracted several quantitative parameters, including total cell count, number of infected cells (Inf), percentage of infected cells, number of intracellular Ot per infected cell (Bac/Inf), and number of perinuclear-localized bacteria per infected cell (NucBac/Inf), to identify candidate host factors influencing various aspects of Ot infection (Fig. 1B). We defined two distinct hit phenotypes corresponding to sequential stages of infection: (1) bacterial entry into host cells and (2) translocation of Ot from the cytosol to the perinuclear region (Fig. 1C). These phenotypes could be selectively enhanced or inhibited by siRNA transfection (Fig. 1D).

During the screen, we observed variability in host cell numbers across siRNA plates due to siRNAs that either promoted cell proliferation or induced cell death (Figs. 1E-F). For example, in plate P37, we categorized siRNAs into three groups (n = 30 per group) based on final host cell densities relative to the in-plate scramble: lower (blue), similar (green), and higher (red) host cell density (Figs. 1F-G). When comparing the number of intracellular Ot per infected cell to the in-plate scramble, we found that siRNAs associated with higher host cell density (red) led to decreased Bac/Inf in 9 out of 30 cases, and three siRNAs from the group with similar host cell density (green) significantly inhibited Ot entry. No inhibitory effect on Ot entry was identified among siRNAs in the lower-density group (blue) (Fig. 1H). This finding indicated that host cell density influences Ot infectivity, as higher host cell density results in a lower effective multiplicity of infection (MOI) per cell.

To account for variability in host cell density, we developed a normalization strategy using cell-count-matched scramble controls. By adjusting key parameters, including total host cell number and infection percentage, we generated matched controls for each siRNA across three defined host cell density categories (Fig. 1I). This normalization enabled more accurate comparisons of Ot infectivity (Bac/Inf) and facilitated the identification of additional hit siRNAs, such as SLC35B1 and C4orf6 on plate P37, which significantly altered infection independently of their effects on cell proliferation (Fig. 1J).

We integrated hit genes identified using both the conventional in-plate scramble control and the cell-count–matched scramble control, thereby generating a comprehensive hit list for both Ot entry and translocation phenotypes (Fig. 1K, example hit images shown in Supp. Fig. 1B, and S1 Table, sheet “Screen Result”). Hits were categorized into three levels based on their significance relative to the scramble control: High (Top 100), Medium (Top 200), and Low (Top 400). siRNAs that reduced host cell density to less than one-third of the in-plate scramble were classified as toxic and excluded from further analysis (S1 Table, sheet “Toxic siRNAs”). This genome-wide screen enabled the systematic identification of host factors and pathways regulating the early stages of Ot infection, thus setting the stage for further mechanistic studies.

### Identification of gene networks regulating Ot entry and translocation

To define the host cell pathways that modulate Ot infection, we performed a meta-analysis of our RNAi screen hits using Metascape. We categorized the hits into two groups: genes whose depletion either inhibited (Fig. 2) or enhanced (Fig. 3) Ot entry and/or perinuclear translocation.

**Figure 2:**
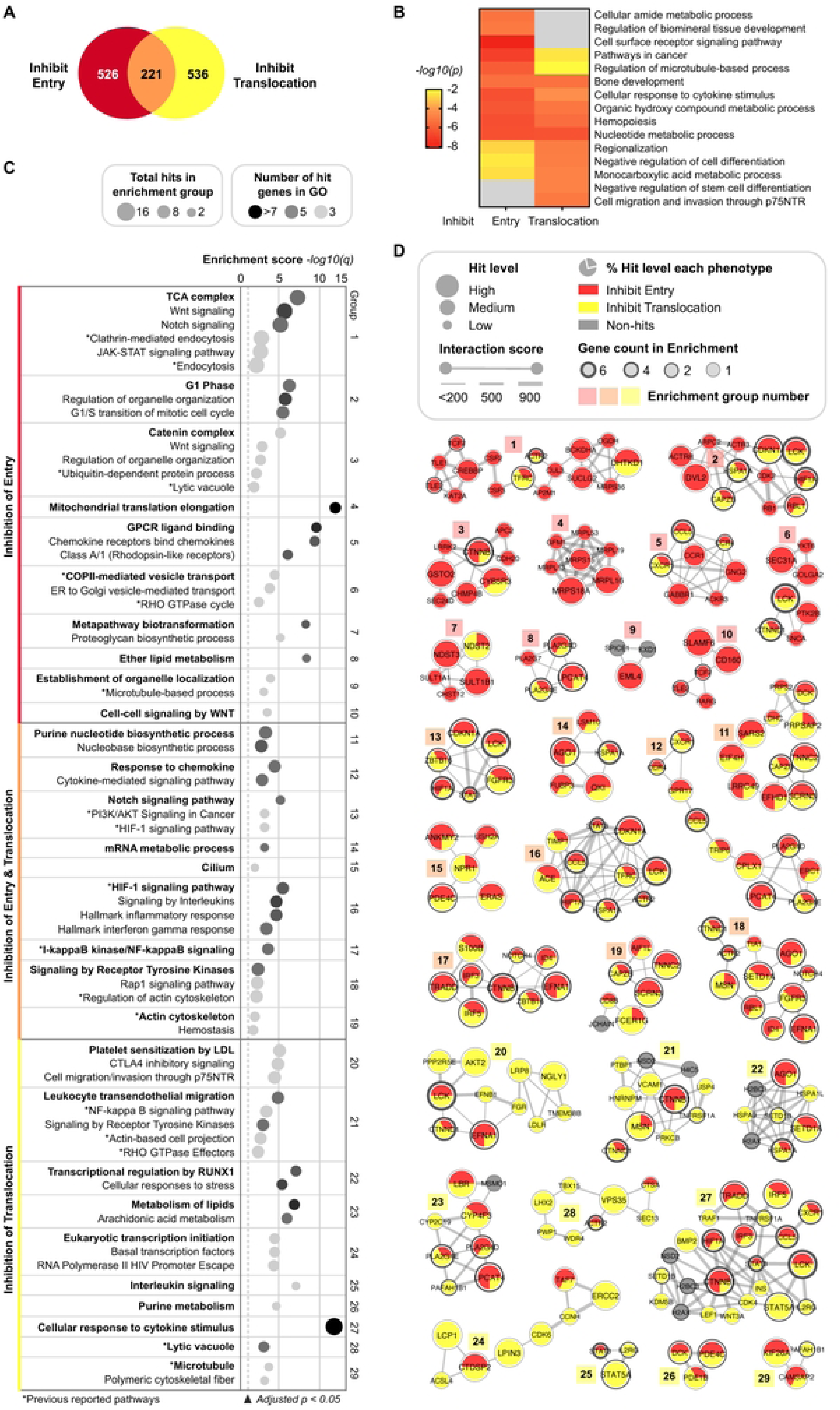
Hit genes and pathways involved in the inhibition of Ot entry and translocation. **A.** Overview of the number of hits causing inhibition of Ot entry, inhibition of Ot translocation, and the intersection of both phenotypes. A total of 1,283 hits were further analyzed using pathway enrichment and protein-protein interaction (PPI) analyses via Metascape. **B.** Top 15 enriched terms for the inhibition phenotypes of Ot entry and translocation. **C-D.** Enriched pathways and the corresponding PPI network of 29 MCODE groups derived from genes involved in the inhibition of Ot entry and translocation. **C.** Twenty-nine significantly enriched pathways (adjusted p-value < 0.05) and their subgroups were identified based on shared genes within the corresponding PPI networks (see panel D). Enrichment scores are presented as -log₁₀(q), where q is the multiple-test adjusted p-value. Each enrichment network was named (in bold) according to a representative term chosen from the top three most significant terms within the subgroup. Superscripts denote previously reported pathways (*). **D.** Protein-protein interaction network of the 29 enrichment groups. Genes are colored according to their inhibitory phenotypes: red for inhibition of Ot entry, orange for inhibition of both Ot entry and translocation, and yellow for inhibition of Ot translocation. Hit genes from the three significance levels (High, Medium, Low) are represented by varying node sizes. Interaction scores, supported by evidence from Metascape, are indicated, and genes identified in multiple groups are highlighted with thicker outlines. The PPI network was annotated and generated using Cytoscape version 3.9.1.

**Figure 3:**
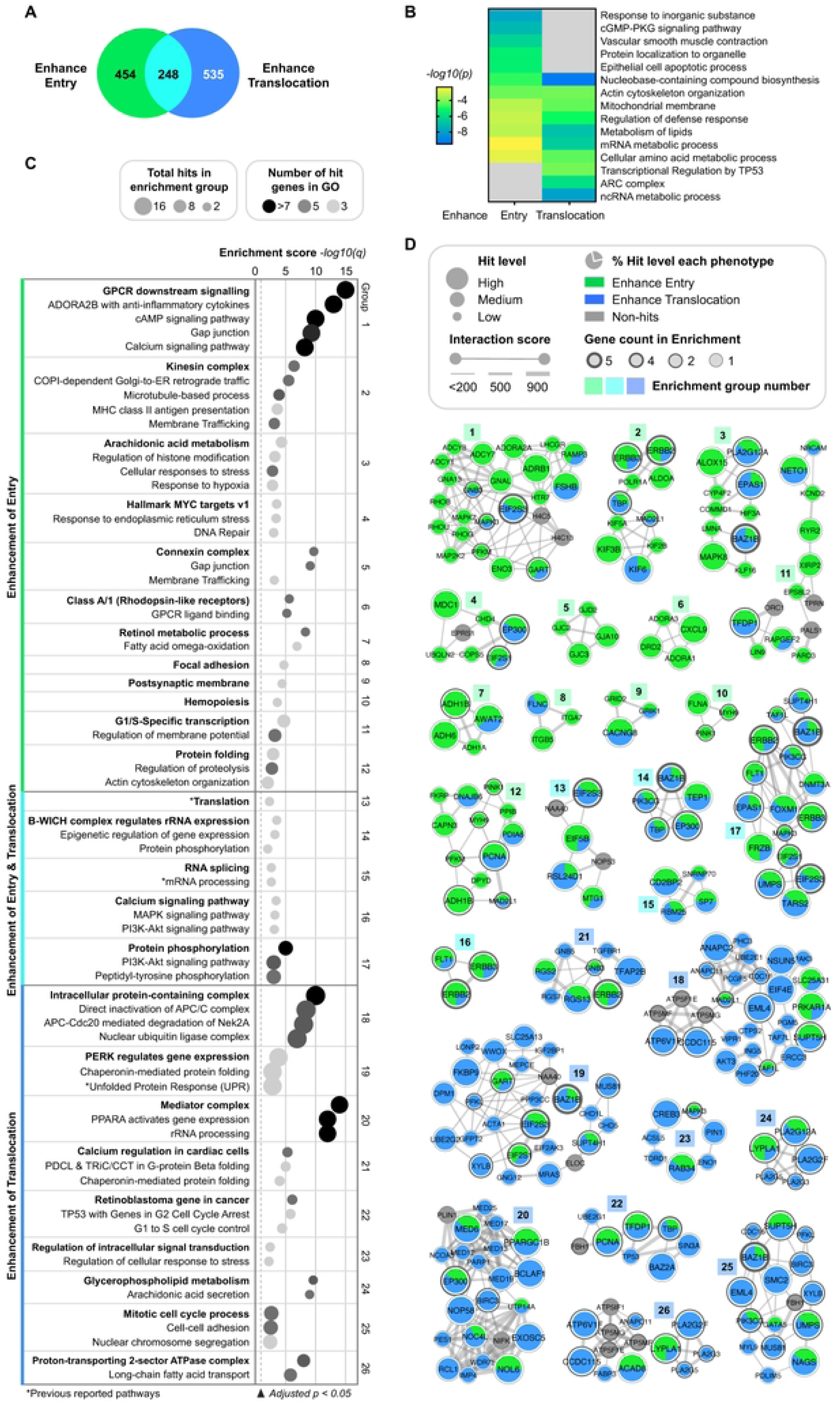
Hit genes and pathways involved in the enhancement of Ot entry and translocation. **A.** Overview of the number of hits causing enhancement of Ot entry, enhancement of Ot translocation, and the intersection of both phenotypes. A total of 1,237 hits were further analyzed using pathway enrichment and protein-protein interaction (PPI) analyses via Metascape. **B.** Top 15 enriched terms for the enhancement phenotypes of Ot entry and translocation. **C-D.** Enriched pathways and the corresponding PPI network of 26 MCODE groups derived from genes involved in the enhancement of Ot entry and translocation. **C.** Twenty-six significantly enriched pathways (adjusted p-value < 0.05) and their subgroups were identified based on shared genes within the corresponding PPI networks (see panel 3D). Enrichment scores are presented as -log₁₀(q), where q is the multiple-test adjusted p-value. Each enrichment network was named (in bold) based on a representative term selected from the top three most significant terms within the subgroup. Superscripts denote previously reported pathways (*). **D.** Protein-protein interaction network of the 26 enrichment groups. Genes are colored according to their enhancement phenotypes: green for enhancement of Ot entry, green-blue for enhancement of both Ot entry and translocation, and blue for enhancement of Ot translocation. Hit genes from the three significance levels (High, Medium, Low) are represented by varying node sizes. Interaction scores, supported by evidence from Metascape, are indicated, and genes identified in multiple groups are highlighted with thicker outlines. The PPI network was annotated and generated using Cytoscape version 3.9.1.

We first analyzed 1,283 siRNA hits that reduced Ot entry and/or translocation (Fig. 2A). Pathway enrichment analysis of these genes revealed the top 15 Gene Ontology (GO) terms (Fig. 2B, S2 Table, sheet “Top100 GO Inhibition”), highlighting pathways involved in cell surface receptor signaling, microtubule-based transport, cytokine responses, and various metabolic processes (e.g., amide, organic hydroxy, nucleotide, and monocarboxylic acid metabolism). These findings suggest that the associated pathways positively regulate Ot infection processes.

To further characterize the biological organization of the inhibitory hits, we applied the MCODE clustering algorithm to the Metascape protein-protein interaction (PPI) network, resulting in 29 modules (Figs. 2C–D, S2 Table, sheet “PPI Inhibition”). Each module was annotated with a representative biological term (bolded) selected from the top three most significantly enriched ontology terms. Asterisks (*) indicate previously reported pathways. Enrichment scores were expressed as –log₁₀(q), where q represents the false discovery rate-adjusted p-value (Fig. 2C). In the PPI network (Fig. 2D), the node color reflects phenotypic categories—red indicates inhibition of Ot entry, orange indicates inhibition of both entry and translocation, and yellow indicates inhibition of translocation alone. Node size corresponds to the significance level of the hit (High, Medium, Low).

Among the key inhibitory modules, Group 1 represented genes involved in clathrin-mediated endocytosis, a well-established pathway critical for Ot entry^13^. Groups 9 and 29 were enriched in microtubule-related genes, consistent with their known role in facilitating perinuclear trafficking of intracellular pathogens^18^. Groups 18, 19, and 21 were associated with actin cytoskeleton regulation, supporting a role for actin remodelling during bacterial internalization. Groups 6 and 21 involved genes related to vesicle transport, which may reflect the process of endosomal escape by intracellular Ot. Additionally, Groups 13 and 16 included components of HIF-1 and PI3K/AKT signaling pathways, both previously implicated in Ot-induced autophagy^15,16^. Genes involved in ubiquitination and vacuolar/lysosomal pathways were also identified (Groups 3 and 28), suggesting engagement of host defense mechanisms^30,31^. Notably, Groups 17 and 21 contained NF-κB signaling components (e.g., IκB), aligning with prior reports that Ot interferes with this immune pathway^32^.

Several newly implicated biological processes were also identified. Group 2 included cell cycle-related genes, while Groups 1, 7, 8, 11, 14, 23, and 26 were enriched for various metabolic pathways. Groups 12, 16, 17, 21, 25, and 27 were associated with immune response and cytokine-mediated signalling pathways. Group 5 highlighted genes involved in G-protein-coupled receptor (GPCR) ligand binding, a pathway not previously linked to Ot infection.

In contrast, analysis of the 1,237 siRNA hits that enhanced Ot entry and/or perinuclear translocation (Fig. 3A) revealed additional insights into negative regulators of infection. GO enrichment analysis of these genes identified the top 15 enriched pathways (Fig. 3B, S2 Table, sheet “Top100 GO Enhance”), including cGMP-PKG signaling, epithelial cell apoptosis, regulation of host defense responses, and diverse metabolic processes involving lipids, mRNA, ncRNA, and amino acids. These pathways appear to restrict Ot infection, and their depletion may promote bacterial entry or early intracellular accumulation.

MCODE clustering of the enhancement-associated genes produced 26 modules (Figs. 3C–D, S2 Table, sheet “PPI Enhance”), with enrichment scores and pathway annotations derived as described above. In the enhancement PPI network (Fig. 3D), nodes were colored according to phenotype: green for enhancement of Ot entry, green-blue for enhancement of both entry and translocation, and blue for enhancement of translocation alone. Node sizes similarly reflected the significance of each hit.

Notable enhancement-associated modules included Group 13, which contained translation-related genes. The suppression of these genes may relieve competition for host-derived amino acids, thus supporting Ot replication. Group 15 involved mRNA processing, suggesting an additional regulatory layer in host biosynthetic activity. Group 19 was enriched for genes associated with the unfolded protein response (UPR), a pathway previously reported to be induced by Ot, along with suppression of ER-associated degradation (ERAD), to enhance amino acid availability^33^. Furthermore, Groups 18, 22, and 25 encompassed cell cycle-related pathways, underscoring their potential role in restricting Ot infection.

Collectively, these analyses provide a comprehensive systems-level map of host gene networks that either promote or limit Ot infection. The identification of both well-characterized and novel regulatory pathways, including those related to cytoskeletal organization, vesicle trafficking, immune signaling, cell cycle control, and cellular metabolism, offers valuable insights into host-pathogen interactions and reveals candidate pathways for future functional validation and therapeutic targeting.

### Validation of primary screen hits using pharmacological inhibitors and a secondary siRNA screen

We next employed pharmacological inhibitors and an independent secondary siRNA screen to validate our primary siRNA screen. First, we selected drugs that target pathways identified to reduce Ot entry or perinuclear translocation in the primary screen (S2 Table, sheet “PPI Inhibition”) and performed a drug screen to assess their effects on these two processes. All compounds were tested at concentrations that minimized cytotoxicity in HeLa cells (see Methods). Overall, 18 drugs targeting 13 pathways were tested. Genes were grouped into the pathways described in Fig. 2C, with selected genes from the primary screen shown as colored bars (red: inhibits only entry; orange: inhibits both entry and translocation; yellow: inhibits only translocation), while drug results are presented in gray (Figs. 4A–M). For pathways affecting Ot entry, fold changes were calculated from the number of intracellular bacteria per infected cell (Bac/Inf; Figs. 4A–F, H, K). For pathways affecting Ot translocation, fold changes were calculated from the number of perinuclear-localized bacteria per infected cell (NucBac/Inf; Figs. 4G, I, J, L, M).

**Figure 4:**
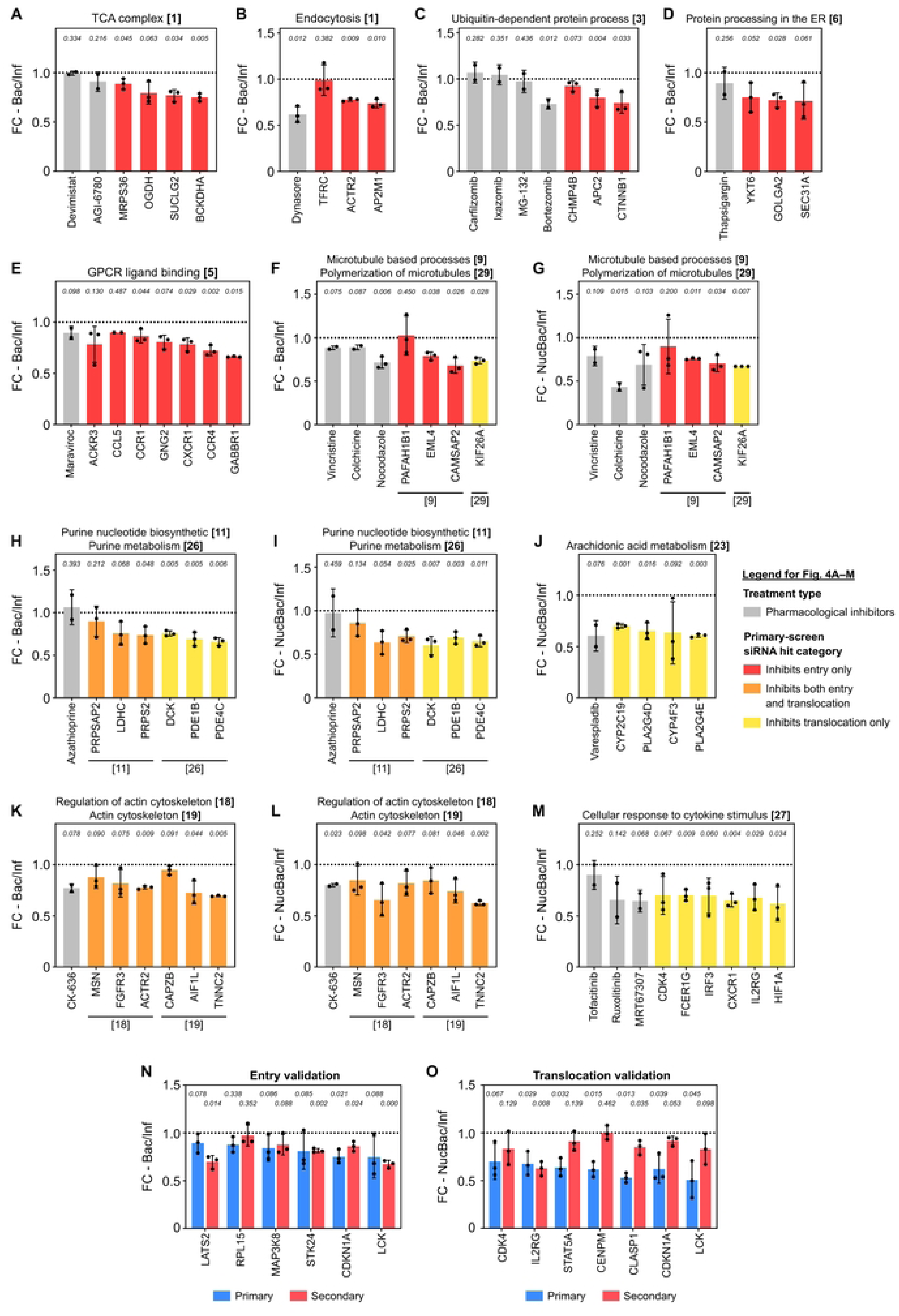
Validation of primary screen hits using pharmacological inhibitors and a secondary siRNA screen. A-M. Pharmacological validation of primary-screen hits affecting Ot entry and translocation. Eighteen drugs targeting 13 pathways identified from the primary screen were tested at non-toxic doses in HeLa cells (Fig. 2C, S2 Table, sheet “PPI Inhibition”; see Methods). Enrichment groups are labeled by group number according to Fig. 2C. Gray bars represent drug-treated conditions, whereas colored bars represent selected siRNA hits from the primary screen shown in Fig. 2D. Red indicates hits that inhibit Ot entry only, orange indicates hits that inhibit both entry and translocation, and yellow indicates hits that inhibit translocation only. Ot entry was quantified as the number of intracellular bacteria per infected cell (Bac/Inf), and Ot translocation was quantified as the number of perinuclear bacteria per infected cell (NucBac/Inf). Fold changes were calculated relative to the corresponding control. Data represent mean ± SD from three experiments. Statistical significance was assessed using Student’s t-test in GraphPad Prism and is indicated above each bar. **A.** Inhibition of Ot entry by Devimistat and AGI-6780 targeting the TCA cycle pathway. **B.** Inhibition of Ot entry by Dynasore, targeting clathrin-mediated endocytosis. **C.** Inhibition of Ot entry by Carfilzomib, Ixazomib, MG-132, and Bortezomib, targeting ubiquitin-dependent protein processing. **D.** Inhibition of Ot entry by Thapsigargin, interfering with protein processing in the ER. **E.** Inhibition of Ot entry by Maraviroc, targeting the GPCR ligand binding. **F-G.** Inhibition of both Ot entry and translocation by drugs targeting microtubule-based processes and polymerization of microtubules: Vincristine, Colchicine, and Nocodazole. **H-I.** Evaluation of Ot entry and translocation after Azathioprine treatment, targeting purine nucleotide biosynthesis and purine metabolism. **J.** Inhibition of Ot translocation by Varespladib, targeting host pathways involved in arachidonic acid metabolism. **K-L.** Inhibition of both Ot entry and translocation via modulation of the actin cytoskeleton using CK-636. **M.** Inhibition of Ot translocation by Tofacitinib, MRT67307, and Ruxolitinib, targeting cellular response to cytokine stimulus pathways. **N.** Secondary siRNA validation of entry hits. Blue bars represent primary-screen results, and red bars represent secondary-screen results obtained using RNAi reagents from an independent supplier. Ot entry was quantified as the fold change in the number of intracellular bacteria per infected cell (FC - Bac/Inf). **O.** Secondary siRNA validation of translocation hits. Blue bars represent primary-screen results, and red bars represent secondary-screen results obtained using RNAi reagents from an independent supplier. Ot translocation was quantified as the fold change in the number of perinuclear bacteria per infected cell (FC - NucBac/Inf). Data are presented as mean ± SD from three independent experiments. Statistical significance compared with controls was determined using Student’s t-test in GraphPad Prism and is indicated above each bar.

Several novel pathways emerged from this analysis and were supported by pharmacological validation. For example, the TCA cycle was significantly enriched among regulators of Ot entry (–log₁₀(q) = 7.3; Fig. 2C), and inhibition with AGI-6780 produced a corresponding reduction in infection (Fig. 4A). Similarly, targeting GPCR signaling with Maraviroc impaired Ot entry, consistent with its high enrichment score (–log₁₀(q) = 9.6; Fig. 4E).

In parallel, previously established host pathways showed concordant effects. Inhibition of endocytosis (Dynasore), ubiquitin-mediated protein processing (Bortezomib), and ER function (Thapsigargin) all reduced Ot infection, validating the screening approach (Figs. 4B–D). Disruption of cytoskeletal systems further confirmed their central role, as microtubule inhibitors (Vincristine, Colchicine, Nocodazole) and actin modulation (CK-636) impaired both bacterial entry and translocation (Figs. 4F–G, 4K–L).

Additional pathways, including arachidonic acid metabolism and cytokine signaling, were specifically associated with bacterial translocation, as inhibition with Varespladib, Ruxolitinib, and MRT67307 reduced perinuclear localization of Ot (Figs. 4J, 4M). In contrast, inhibition of purine metabolism with Azathioprine did not significantly affect infection under these conditions (Figs. 4H–I), suggesting context-dependent roles for this pathway.

To validate the primary screen, we performed a secondary siRNA screen using independent RNAi reagents from a different vendor. Efficient target knockdown was confirmed at 48 and 72 hours post-transfection (Supp. Fig. 1C). We then assessed 11 candidate genes identified in the primary screen as regulators of Ot entry or translocation. Knockdown of selected genes reproduced the corresponding primary-screen phenotypes, reducing Ot entry and/or perinuclear translocation in the secondary screen (Figs. 4N–O), supporting the reproducibility and robustness of the primary screening results.

### Differential effects of cell cycle gene knockdown on Ot entry and localization

Our siRNA screen revealed that genes involved in host cell cycle regulation exert distinct and, in some cases, opposing effects on Ot entry and intracellular localization. Knockdown of genes associated with the G1 phase (Group 2; Fig. 2C–D), including CDK2, CDKN1A, and RB1, reduced Ot entry. In contrast, depletion of genes involved in G1/S-specific transcription (Group 11; Fig. 3C–D), such as TFDP1 and NETO1, as well as genes regulating the G2 phase (Group 22; Fig. 3C–D), including PCNA and TP53, enhanced Ot entry and perinuclear translocation. These opposing effects indicate that host cell cycle phase influences susceptibility to infection and subsequent bacterial trafficking, suggesting that distinct cell cycle states provide intracellular environments that differentially restrict or support Ot infection.

In the primary screen, siRNA knockdown was performed 48 hours prior to Ot infection, a timeframe during which perturbation of cell cycle regulators could alter the baseline cellular state encountered by the pathogen. Previous studies have shown that depletion of specific cell cycle genes induces arrest at defined checkpoints (Fig. 5A). For example, CDK4 knockdown promotes G1 arrest^34^, and CDK6 targeting inhibits RB phosphorylation and induces G1 arrest^35^. More broadly, the CDK–RB–E2F pathway regulates G1/S progression, and loss or knockdown of RB1 disrupts cell-cycle control by affecting G1/S and G2/M transitions and promoting inappropriate S-phase entry^36–38^. PCNA knockdown has been reported to cause S-phase arrest^39^. In addition, SMC2 downregulation suppresses WNT-activated cell proliferation and has been proposed as a therapeutic target, whereas knockdown of mutant TP53 induces G2-phase arrest and apoptosis^40,41^.

**Fig. 5:**
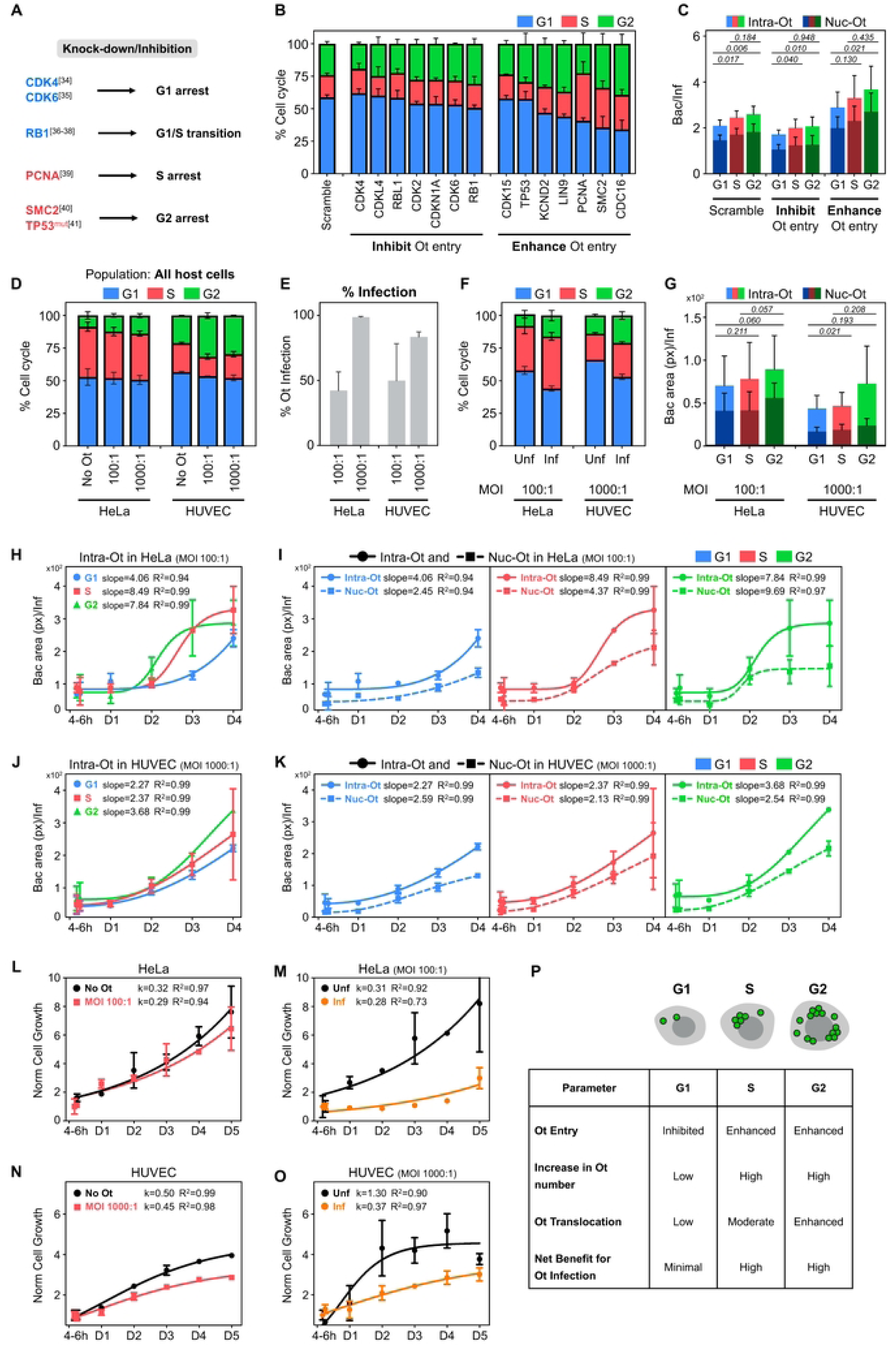
Cell cycle-dependent regulation of Ot infection and host cell proliferation. **A.** Schematic showing the known effects of knockdown or targeting of cell cycle-related genes on cell cycle arrest in specific phases based on previous studies: G1 arrest with CDK4 knockdown and CDK6 targeting, altered G1/S progression with RB1 loss or knockdown, S-phase arrest with PCNA knockdown, and G2-phase arrest or growth arrest/apoptosis with SMC2 downregulation and mutant TP53 knockdown (TP53^mut^). **B.** Distribution of cell cycle phases in HeLa cells following knockdown of selected cell cycle genes retrieved from the primary screen: seven hit genes from Group 2 (inhibiting Ot entry) and seven hit genes from Groups 11 and 22 (enhancing Ot entry and translocation), compared to scramble control. No clear overall pattern was observed between inhibitory and enhancing hits. **C.** Quantification of intracellular Ot (Intra-Ot) and perinuclear Ot (Nuc-Ot) in HeLa cells across cell cycle phases for the groups of hit genes shown in panel 5B, revealing significantly higher bacterial loads in S and G2 phases compared to G1. Statistical significance was determined using the Mann–Whitney U test **D-G.** Early effects of Ot infection on host cell cycle distribution in HeLa and HUVEC cells. Cells were infected for 4-6 hours at varying MOIs and labeled with EdU and Hoechst to determine cell cycle phases. **D.** Distribution of cell cycle phases in HeLa and HUVEC cells at 4-6 hours post-infection with Ot. **E.** Percent infection in HeLa and HUVEC cells at 4-6 hours post-infection with Ot. **F.** Distribution of cell cycle phases in infected cells and uninfected cells of HeLa (MOI 100:1) and HUVEC (MOI 1000:1) at 4-6 hours post-infection. Infected populations in both cell types showed a significant increase in S and G2 phases relative to uninfected cells. **G.** Quantification of intracellular Ot (Intra-Ot) and perinuclear Ot (Nuc-Ot) area retrieved from infected cells of panel 5F across cell cycle phases, revealing that S and G2 cells harbor higher bacterial loads than G1 cells. Statistical significance was determined using Student’s t-test. **H-K.** Long-term analysis of Ot colony expansion in HeLa (MOI 100:1) and HUVEC (MOI 1000:1) cells across cell cycle phases. Microcolony areas in G1, S, and G2 cells were measured at 4-6 hours, 1, 2, 3, and 4 days post-infection and fitted with a sigmoidal 4-parameter logistic model. The hill slope represents the relative Ot expansion rate. **H.** Ot expansion in HeLa cells. **I.** Comparison of intracellular Ot (Intra-Ot) versus perinuclear Ot (Nuc-Ot) expansion in HeLa cells. **J.** Ot expansion in HUVEC cells. **K.** Comparison of Intra-Ot versus Nuc-Ot expansion in HUVEC cells. **L-O.** Analysis of host cell proliferation under Ot infection. Host cell numbers were monitored from 4 hours to 5 days post-infection, and proliferation rate constants (k) were determined using exponential models (Malthusian for HeLa; Gompertz for HUVEC). **L.** Comparison of HeLa cell proliferation in uninfected (No Ot) versus Ot-infected (MOI 100:1) conditions. **M.** Comparison of proliferation between uninfected (Unf) and infected (Inf) HeLa cells within the same culture (MOI 100:1, ∼40% infection at 4-6 hours). **N.** Comparison of HUVEC cell proliferation in uninfected (No Ot) versus Ot-infected (MOI 1000:1) conditions. **O.** Comparison of proliferation between uninfected (Unf) and infected (Inf) HUVEC cells within the same culture (MOI 1000:1, ∼80% infection at 4-6 hours). **P.** Schematic diagram and accompanying summary table integrating our findings on how host cell cycle phases influence Ot infection. The diagram illustrates differences between G1, S, and G2 phases regarding Ot entry, increase in Ot number, Ot translocation, and the overall net benefit for Ot infection.

To assess how these perturbations influence host cell cycle distribution in our system, we quantified DNA content using integrated Hoechst intensity from high-content imaging data and classified individual cells into G1, S, and G2 phases. Mitotic cells were excluded from analysis because the absence of an intact nuclear envelope prevented reliable measurement of perinuclear Ot. Therefore, the 4N population analyzed here is referred to as G2 rather than G2/M. We analyzed representative genes from Group 2 (G1-associated; inhibitory for Ot entry) and compared them with genes from Group 11 (G1/S transcription) and Group 22 (G2 regulation), which enhanced Ot entry and/or translocation. Although these perturbations altered cell cycle distributions (Fig. 5B), no consistent pattern distinguished genes that inhibited versus promoted infection.

These findings suggest that cell cycle redistribution alone does not fully explain the observed infection phenotypes. Instead, we hypothesized that the intrinsic cell cycle state at the time of infection influences susceptibility to Ot. Supporting this, spatial analysis revealed that intracellular bacteria (Intra-Ot) and perinuclear-localized bacteria (Nuc-Ot) were present across all phases, but were significantly more abundant in S and G2 cells compared to G1 (Fig. 5C). This distribution indicates that S and G2 phases provide a more permissive environment for Ot entry and early intracellular accumulation.

### Ot preferentially accumulates in S and G2 and is associated with host cell cycle delay during early infection

To investigate the effect of Ot infection on host cell cycle progression, we infected HeLa and HUVEC cells at varying multiplicities of infection (MOIs) and quantified bacterial localization and cell cycle status at 4–6 hours post-infection (Fig. 5D–G). Cell cycle phase was determined by EdU incorporation and Hoechst staining, enabling classification of individual cells into G1, S, and G2 phases (Supp. Fig. 2A). After 4-6 hours of Ot infection, HeLa cells showed a slight increase in the proportion of cells in the G2 phase, while HUVEC cells showed a more pronounced enrichment in G2 at higher MOIs (100:1 and 1000:1) (Fig. 5D). To distinguish effects within mixed populations, we analyzed infected and uninfected cells separately at MOI 100:1 (HeLa, ∼40% infection) and MOI 1000:1 (HUVEC, ∼80% infection) (Fig. 5E). In both cell types, infected cells were significantly enriched in S and G2 phases compared to uninfected cells (Fig. 5F).

These results suggest that, rather than accelerating overall cell cycle progression, Ot infection is associated with a shift toward, and potential delay within, S and G2 phases at early time points. Consistent with this, intracellular bacterial signal, measured as Intra-Ot area, was higher in S and G2 cells than in G1, while perinuclear localization, measured as Nuc-Ot area, showed a modest increase in G2 cells (Fig. 5G). Together, these findings indicate that S and G2 phases provide a more permissive environment for early bacterial accumulation and suggest that Ot infection may influence host cell cycle dynamics to favor these states.

### Pathogen-induced S and G2 phase delays support enhanced Ot colony expansion

Having established that early Ot infection is associated with enrichment of host cells in S and G2 phases, we next examined how these cell cycle states influence intracellular bacterial expansion. We quantified Ot microcolony area in G1, S, and G2 cells of HeLa (MOI 100:1) and HUVEC (MOI 1000:1) cells over time (4–6 hours to 4 days post-infection) (Fig. 5H–K; Supp. Fig. 2B–E).

In HeLa cells, Ot colonies in S and G2 phases exhibited markedly higher expansion rates than those in G1, with hill slopes of 8.49 and 7.84 compared to 4.06, respectively (Fig. 5H). During early time points (4–6 hours and day 1), bacterial area remained relatively stable, consistent with a lag phase of intracellular establishment. From day 2 onwards, colonies in S and G2 cells transitioned into rapid expansion, indicating that these phases support more efficient bacterial growth.

Analysis of subcellular localization revealed that perinuclear-associated bacteria (Nuc-Ot) expanded more slowly than total intracellular bacteria (Intra-Ot) across all cell cycle phases (Fig. 5I). Quantitatively, Nuc-Ot expansion rates (hill slopes) were 2.45, 4.37, and 9.69 in G1, S, and G2 phases, respectively, and were consistently lower than corresponding Intra-Ot expansion rates. At day 2, Nuc-Ot and Intra-Ot areas were comparable in S and G2 cells, indicating transient accumulation of bacteria in the perinuclear region. At later time points (days 3–4), perinuclear expansion lagged behind total intracellular growth, consistent with redistribution of bacteria into the cytosol prior to egress.

In HUVEC cells, overall expansion rates were lower than in HeLa cells, with hill slopes of 2.27, 2.37, and 3.68 in G1, S, and G2 phases, respectively (Fig. 5J). Temporal transitions were less pronounced, and no clear exponential phase was observed. Nevertheless, colonies in S and G2 phases consistently expanded more than those in G1, indicating that the association between cell cycle state and bacterial growth is conserved across cell types. Analysis of subcellular localization showed that Nuc-Ot expansion remained slightly lower than Intra-Ot, with rates of 2.59, 2.13, and 2.54 in G1, S, and G2 phases, respectively (Fig. 5K). The difference between perinuclear and total intracellular populations was most evident in G2 cells at day 2, after which bacterial expansion increasingly occurred in non-perinuclear regions, consistent with redistribution prior to exit.

Together, these findings demonstrate that S and G2 phases provide a more permissive environment for Ot expansion and suggest that enrichment in these states enhances intracellular population growth while coordinating spatial redistribution during later stages of infection.

### Ot infection impairs overall host cell proliferation to sustain its replicative niche

Having established that Ot preferentially accumulates in S and G2 phase cells, and that infection is associated with enrichment of host cells in S and G2 phases at early stages of infection, we next examined whether these changes are accompanied by altered host cell proliferation over longer time scales. HeLa (MOI 100:1) and HUVEC (MOI 1000:1) cells were monitored from 4 hours to 5 days post-infection, and proliferation dynamics were modeled using exponential (Malthusian) growth for HeLa cells and a Gompertz model for HUVEC cells. Proliferation rate constants (k) were used to quantify host cell division kinetics.

In HeLa cells, uninfected controls exhibited a proliferation rate of k = 0.32, whereas Ot-infected populations showed a reduced rate of k = 0.29 (Fig. 5L). Within mixed populations (∼40% infection at 4–6 hours; Fig. 5E), infected cells proliferated more slowly than uninfected cells (k = 0.28 vs 0.31; Fig. 5M), indicating a direct association between infection and reduced host cell division. The increase in infected cell number over time reflects both proliferation of infected cells and secondary infection of previously uninfected cells, which cannot be fully separated in this assay.

A similar but more pronounced effect was observed in HUVEC cells. Uninfected populations exhibited a proliferation rate of k = 0.50, whereas infected populations showed a reduced rate of k = 0.45 (Fig. 5N). In mixed cultures with ∼80% initial infection (Fig. 5E), infected cells displayed substantially lower proliferation (k = 0.37), while the remaining uninfected cells exhibited higher apparent growth (k = 1.30; Fig. 5O), consistent with expansion of the uninfected subpopulation. Together, these findings indicate that Ot infection is associated with reduced host cell proliferation across cell types. Taken together with the enrichment of S- and G2-phase cells and enhanced bacterial expansion in these states, these results support a model in which Ot preferentially accumulates in specific host cell cycle phases while limiting host cell division, thereby stabilizing an intracellular environment conducive to sustained bacterial growth and subsequent dissemination.

## Discussion

This work has led to the identification of extensive networks of host cell genes that orchestrate the early stages of infection by the obligate intracellular bacterium Ot. In addition to secondary validation of hits via both siRNA and pharmacological inhibition, our findings are reinforced by the recovery of numerous pathways previously implicated in Ot infection—such as clathrin-mediated endocytosis, actin cytoskeleton dynamics, microtubule-based processes, surface proteoglycans, the unfolded protein response, and NF-κB signaling. Moreover, the genome-wide RNAi screen revealed novel pathways not previously linked to Ot entry and translocation, including those involved in host cell metabolism, cytokine response, and host cell cycle regulation, thereby generating new testable hypotheses regarding Ot infection.

Analysis of the phenotypic screen data presented significant challenges, particularly due to variability in host cell numbers following siRNA transfection. Variations in total host cell counts can alter the Ot-to-host cell ratio, complicating the interpretation of gene knockdown effects on Ot infection phenotypes. Initially, our standard scramble control (in-plate scramble) was insufficient for accurately normalizing these differences, which meant that only genes with extreme phenotypes were reliably identified. By developing a novel cell-count-matched scramble control that pairs each siRNA condition with an equivalent host cell density, we were able to address this variability and recover a more comprehensive and accurate set of hit genes. This refined approach not only enhanced the reliability of our findings but also underscored the importance of considering host cell proliferation dynamics when dissecting complex host-pathogen interactions.

Our results reveal a critical interplay between Ot infection and host cell cycle dynamics that unfolds in several distinct phases (see Fig. 5P). First, our screening and knockdown experiments demonstrate that Ot preferentially accumulates in host cells in the S and G2 phases rather than in G1. In HeLa cells, inhibition of G1 phase genes reduced Ot entry, whereas knockdown of genes governing the G1/S transition and G2 phase enhanced both entry and translocation. This observation suggests that the intracellular environment during S and G2, possibly characterized by increased metabolic activity and elevated levels of nucleotides and amino acids, offers optimal conditions for Ot entry and colony expansion. Factors such as variations in lipid composition, endocytic activity, or surface receptor abundance might also contribute. Overall, our data strongly indicate that the cell cycle stage itself plays a pivotal role. This strategy of exploiting specific cell cycle phases is shared by other intracellular pathogens; for instance, *Listeria monocytogenes* induces an S-phase delay to favor bacterial propagation^42^, and *Staphylococcus aureus* delays the G2/M transition to enhance infective efficiency and intracellular accumulation^43^.

Second, temporal population dynamics analyses reinforce these findings. In HeLa cells, Ot colonies in S and G2 phases rapidly transitioned into an exponential expansion phase following an initial lag period, exhibiting significantly higher relative expansion rates than those observed in G1. A similar, albeit less pronounced, trend was observed in HUVEC cells. In both cell types, perinuclear localization of Ot was initially constrained, with S-phase cells supporting only moderate transition to non-perinuclear cytoplasmic localization compared with G2-phase cells.

This suggests that bacterial maturation and exit are more likely to occur in S/G2 phase cells compared with G1.

Third, we observed that Ot infection also impacts host cell proliferation. Comparison of the proliferation rate constants (k values) of infected versus uninfected cells revealed that Ot-infected populations in both HeLa and HUVEC cells proliferate more slowly. This reduction in host cell division may create a more stable intracellular environment that favors bacterial colony expansion by limiting competition for essential resources and reducing the disruptive effects of rapid cell turnover. The underlying mechanism might involve secreted cyclomodulin-type effector proteins, although further investigation is needed.

Collectively, these findings illustrate that Ot not only benefits from the favorable metabolic conditions of the S and G2 phases for accelerated colony expansion but may also influence host cell cycle progression and proliferation. Our phenotypic observations strongly align with recent mechanistic discoveries demonstrating that a different strain of Ot, Ikeda, actively arrests the host cell cycle at the S phase. This manipulation is orchestrated by the bacterial nucleomodulin Ank13, which represses *TP53* transcription to deplete p53 levels, thereby overriding canonical cell cycle checkpoints and preventing early host cell apoptosis to secure a stable replicative niche^21^. This dual strategy likely enables Ot to establish a stable niche and optimize its intracellular accumulation and dissemination. The summary diagram in Fig. 5P encapsulates these observations. Future studies focusing on the specific cell cycle regulatory pathways and host signaling events affected by Ot will be critical to further elucidate the mechanisms underlying these observations.

There are limitations associated with our study. Redundancy in host gene networks means that some genes involved in Ot infection will not cause a phenotype following single-gene depletion. Similarly, the use of a knockdown rather than a knockout screen means that residual protein activity may suffice to perform the required function, resulting in false-negative hits. Another limitation of our screen is that although we selected a time point for fixation prior to the expected onset of massive bacterial accumulation, we cannot distinguish between an inhibitory phenotype being caused by a decrease in bacterial entry or an increase in bacterial killing by host cells, nor can we distinguish between an enhancement phenotype being caused by an increase in bacterial entry or early bacterial replication. Further experiments will be needed to distinguish modes of action of specific hits. Finally, our screen used the human epithelial cell line HeLa and the Ot clinical isolate UT76. Whilst many of the pathways uncovered by our screen are likely to be universal, it is possible that some pathways would differ in different cell lines or with different Ot strains.

We recently showed that Ot differentiates into two forms during its intracellular infection cycle, a metabolically active intracellular form and a metabolically inactive extracellular form^14^. We showed that these forms enter host cells using different mechanisms, although both were competent for subsequent intracellular colony expansion. The current screen was carried out using bacteria harvested from the intracellular fraction, and it is expected that there may be some differences in the pathways exploited by the extracellular form, which could be explored in future studies.

This is the first genome-wide screen for host cell genes involved in infection of any Rickettsiales, and we hope this will lead to new discoveries based on the hits identified here, as well as providing a template for similar analyses in related obligate intracellular bacteria.

## Materials and Methods

### Cells lines, strains, and growth conditions used in this work

HeLa was generously gifted from the Mitchison lab at Harvard University, USA. HeLa cells were cultured in DMEM supplemented with 10% FBS. HUVEC cell was cultured and maintained in Medium 200 supplemented with LVES. Both cell types were incubated at 37 °C, 5% CO_2_, and 95% air.

### Genome-Wide RNAi Screening

We performed the genome-wide siRNA screen using the Silencer Select siRNA - Human Whole Genome library (Ambion), which targets 20,655 genes with pools of three distinct oligonucleotides per gene. siRNAs were pre-spotted by the Centre for High Throughput Phenomics (CHiP-GIS) at 200 nM in 5 µL RNase-free water into black, clear-bottom 384-well plates (Corning 3712). Plates were thawed at room temperature and briefly spun down prior to use.

For transfection, 5 µL of Opti-MEM Reduced Serum Medium (Thermo Fisher Scientific #31985-070) was dispensed into each well using a Multidrop Combi. In parallel, 0.1 µL Lipofectamine 3000 (Thermo Fisher Scientific #L3000-008) was mixed with 10 µL Opti-MEM Reduced Serum Medium and then dispensed into each well. The plates were incubated at room temperature for 30 minutes to allow complex formation.

Next, 2,000 HeLa cells (passage number ≤20) in 30 µL DMEM supplemented with 10% FBS were added to each well, yielding a final siRNA concentration of 20 nM in media containing 5% serum. The cells were incubated at 37 °C in 5% CO_2_ for 24 hours. Following this, the transfection media was removed manually, and 50 µL of fresh DMEM with 5% FBS was added. The cells were further incubated for an additional 24 hours prior to infection.

For infection, Ot strain UT76 was added in 25 µL DMEM with 5% FBS at a multiplicity of infection (MOI) of 200:1 (yielding an actual MOI of 3:1 based on image analysis from in-plate scramble controls). To synchronize infection, the plates were centrifuged at 700 x g for 10 minutes at 25 °C. After incubation at 35 °C in 5% CO_2_ for 3 hours, the Ot-containing medium was removed and replaced with 100 µL fresh DMEM with 5% FBS. The cells were then cultured in a moisture box at 35 °C in 5% CO_2_ for an additional 27 hours.

Finally, cells were fixed with 4% formaldehyde at room temperature for 10 minutes, permeabilized with absolute methanol on ice for 20 minutes, and subsequently labeled with Hoechst (nuclei), a FITC-conjugated antibody against Ot UT76 protein, and Alexa Flour 647 (Evans Blue for cell outlines). Plates were imaged using the Operetta High-Content Imaging System (PerkinElmer).

### High-Content Imaging and Processing

384-well siRNA plates, labeled with fluorescent markers, were imaged using a PerkinElmer Operetta HCS system equipped with a 40x WD objective. The system, integrated with Plate::Handler™, allowed automated imaging of up to 6-9 plates concurrently. For each well, twelve fields were acquired using three fluorescence channels: Hoechst (nuclear stain), FITC (detecting Ot UT76 protein), and Alexa Fluor 647 (cell outline via Evans Blue).

Image segmentation was performed with PerkinElmer’s Columbus software. Nuclear areas were identified using a modified “Method B” based on the Hoechst channel, while whole-cell areas were delineated using an adjusted “Method A” based on the Alexa Fluor 647 channel. Bacterial spots were segmented using an adapted “Method A” from the FITC channel. For each well, Columbus calculated the average number and intensity of identified objects in the Hoechst, FITC, and Alexa Fluor 647 channels.

Several Columbus modules were employed for illumination correction, object removal, and identification of intracellular Ot, as well as Ot adjacent to host nuclei (perinuclear bacteria, NucBac). Artifacts, including edge-touching cells and nuclear objects corresponding to metaphase cells, which lack clear nuclear membranes, were excluded based on nuclear morphology and fluorescence intensity. Mitotic cells were excluded using the Hoechst channel because condensed mitotic chromosomes and nuclear envelope breakdown prevented reliable nuclear segmentation and downstream quantification of perinuclear Ot. We defined infected cells as those containing at least one intracellular bacterium within the whole-cell region. The perinuclear region was defined as a 5-pixel-wide area surrounding the segmented nucleus and was used to quantify perinuclear Ot (NucBac).

### Data Analysis and Hit Identification

Hit gene identification proceeded in three steps. First, siRNAs that reduced host cell density below one-third of the mean scramble control in the same plate were deemed toxic and excluded, eliminating 1,192 siRNAs (S1 Table, sheet “Toxic siRNAs”) and leaving 19,463 non-toxic siRNAs (S1 Table, sheet “Screen Result”).

Second, Ot infectivity for each siRNA was compared to two negative controls: the in-plate scramble and a cell-count-matched scramble. The matched scrambles were generated by adjusting for host cell density factors (total cell number and percentage of infected cells), allowing accurate comparisons of key parameters, namely the number of intracellular Ot per infected cell (Bac/Inf) and the number of bacteria adjacent to the nucleus per infected cell (NucBac/Inf).

In the final step, the entire dataset was analyzed twice independently, using in-plate and matched scrambles, by calculating the Strictly Standardized Mean Difference (SSMD) score for each siRNA relative to its control. Hit genes were classified into three categories, High (Top 100), Medium (Top 200), and Low (Top 400), based on local SSMD thresholding within each siRNA plate. Local thresholds were determined using the median SSMD ± a constant times the interquartile range (S3 Table, sheet “Local SSMD Threshold”). siRNAs with SSMD scores above the threshold were considered hits that enhance Ot infection, whereas those below were considered inhibitory. A comprehensive hit list was then compiled, categorizing hits into four phenotypes: enhancement or inhibition of Ot entry and enhancement or inhibition of Ot translocation. The number of hits per siRNA plate is provided in S3 Table, sheet “Number of Hits in siRNA Plate”.

### Generation of Cell-Count Matched Scramble Controls

To account for variability in host cell density, we generated cell-count matched scramble controls tailored to each phenotype (Ot entry and translocation). First, in-plate scramble data were divided into ten batches based on their time-sequence order during screening. For each batch, we established relationships between selected parameters, using both linear and non-linear models (including polynomial degree 2, exponential, and reciprocal functions) to predict phenotype-specific values.

For Ot entry, six parameters from the in-plate scrambles were used: three biological parameters (total host cell count, number of infected cells, and number of uninfected cells) and three experimental parameters corresponding to replicate order (replicate 1, 2, and 3). These were used to predict the Bac/Inf values for each siRNA. For Ot translocation, an additional parameter (intracellular Ot, Bac/Inf) was incorporated to generate predicted NucBac/Inf values. Seven parent functions were formulated to capture the parameter relationships within each batch. The optimal parent function for each batch was selected based on the minimum L2-norm value (Euclidean norm; S3 Table, sheet “Selected Parent Functions”, labeled in green). A 10-fold cross-validation was then performed on a training dataset to identify the equation with the highest R2 value (S3 Table, sheet “R-square of 10-Fold Cross Validation”). The best equation from each batch was applied to individual siRNA data to accurately predict Bac/Inf and NucBac/Inf values for the matched scrambles. The same approach, with adjusted parameters and parent functions, was used for the Ot translocation analysis.

### Pathway Enrichment Analysis and Protein-Protein Interaction

To visualize enriched pathways associated with sequential Ot infection processes,namely, Ot entry and subsequent translocation from the cytosol to the perinuclear region, ranked gene lists from each phenotype were submitted to the Metascape online tool (https://metascape.org/gp/index.html#/main/step1). Separate meta-analyses were conducted for genes whose knockdown inhibited or enhanced Ot entry/translocation, allowing comparison of enriched terms unique to entry/translocation versus those shared between them. Enrichment analyses were performed across biological processes, canonical pathways, Reactome, and KEGG, yielding the top 100 significantly enriched terms (see S2 Table, sheets “Top100 GO Inhibition” and “Top100 GO Enhance”).

We used Metascape to construct a protein-protein interaction (PPI) network from the unique and merged hit lists for both inhibition and enhancement phenotypes of Ot entry and translocation inhibition. Data from physical core and combined core interactions were integrated, and networks with a minimum size of three genes were retained for analysis. The MCODE algorithm was applied to extract protein interaction complexes from these large networks, and each MCODE cluster was annotated with a representative GO term chosen from the top three most significant terms. The resulting PPI networks and their enrichment scores for both inhibition and enhancement phenotypes are provided in S2 Table, sheets “PPI Inhibition” and “PPI Enhance”. All PPI networks were annotated and generated using Cytoscape version 3.9.1.

### Cell Cycle Analysis

To explore the combined effect of siRNA knockdown and Ot infection on host cell cycle progression in the primary screen, integrated Hoechst fluorescence intensity was quantified for each cell and log₂-transformed. Histograms of these intensity values were generated from control wells, and a Gaussian Mixture Model was applied to identify the two primary peaks corresponding to 2N (G1) and 4N (G2) DNA content. Cutoff values were determined using a bias factor of 1.2 to establish four boundaries. The first and second boundaries were calculated as the mean of the G1 peak minus and plus 1.2 times its standard deviation. Similarly, the third and fourth boundaries were computed as the mean of the G2 peak minus and plus 1.2 times its standard deviation. Cells with intensity values between these two peaks were classified as being in S phase. Individual cells were classified into G1, S, or G2/M phases based on these cutoffs, and the percentage of cells in each phase was calculated for each well. Cells with mitotic morphology were excluded from analysis because nuclear envelope breakdown during M phase prevented reliable segmentation of the nucleus and quantification of perinuclear Ot (NucBac/Nuc-Ot). Therefore, cells with 4N DNA content retained in the analysis were classified as G2 rather than G2/M.

To assess the effect of Ot infection on host cell cycle progression, HeLa and HUVEC cells were infected at varying MOIs and analyzed at 4-6 hours, 1, 2, 3, 4, and 5 days post-infection. Cells were labeled with EdU to mark replicating DNA and with Hoechst to stain total DNA, allowing classification of cell cycle phases (G1, S, or G2/M).

For EdU labeling, cells were treated with 10 µM EdU for 1 hour and then fixed in 4% PFA for 10 minutes. After permeabilization with cold methanol, the EdU reaction was performed using a freshly prepared mixture of 0.1 M Tris (pH 8.5), 10 µM carboxyrhodamine-110 azide dye, 4 mM CuSO_4_, and 50 mM ascorbic acid, incubated for 1 hour in the dark, and washed three times with 0.1% PBS-T. Cells were then stained with Hoechst for nuclei and with a FITC-conjugated antibody against Ot UT76 protein. Imaging was conducted on a confocal microscope and processed using PerkinElmer’s Columbus software.

For cell cycle analysis, at least 2,000 cells per condition were plotted as a scatter between integrated Hoechst intensity (reflecting DNA content) and integrated EdU intensity (indicating DNA synthesis). Using the bimodal distribution in each axis, cutoffs were determined to distinguish negative from positive staining. A Gaussian Mixture Model (GMM) was then applied to cluster cells into three segments: G1 phase (low Hoechst and low EdU), S phase (positive EdU), and G2/M phase (high Hoechst and low EdU).

### Analysis of Ot Colony Expansion and Host Cell Proliferation

For analysis of Ot colony expansion, HeLa cells (infected at an MOI of 100:1) and HUVEC cells (infected at an MOI of 1000:1) were cultured and sampled at 4-6 hours, 1 day, 2 days, 3 days, and 4 days post-infection. At each time point, images were acquired using high-content imaging, and cells were segmented based on cell cycle phase (G1, S, and G2/M) as previously described. In each phase, Ot microcolony area was quantified by measuring the total intracellular Ot fluorescent area per infected cell. These measurements were fitted to a sigmoidal 4-parameter logistic (4PL) model, with the resulting hill slope representing the relative Ot expansion rate. In addition, intracellular Ot (Intra-Ot) and perinuclear Ot (Nuc-Ot) areas were compared to assess differences in bacterial localization.

Host cell proliferation was simultaneously monitored from 4 hours to 5 days post-infection. Normalized total cell numbers were quantified using automated image segmentation, and proliferation rate constants (k) were determined by fitting the data to exponential growth models, using a Malthusian model for HeLa cells and a Gompertz model for HUVEC cells. Comparisons were made between uninfected (No Ot) and Ot-infected conditions, as well as between infected and uninfected populations within the same culture (approximately 40% infection in HeLa and 80% infection in HUVEC at 4-6 hours).

### Pharmacological Inhibition Assays

Eighteen drugs targeting 13 pathways (as identified in Fig. 2C-D and validated in Fig. 4) were selected for evaluation. Eight thousand HeLa cells were seeded per well in 8-well iBidi-treated slides using DMEM supplemented with 5% FBS. For most drugs, treatments were initiated 24 hours prior to infection at the indicated concentrations, while colchicine, dynasore, nocodazole, and vincristine were added 1 hour before infection. Control wells received the corresponding vehicle solution. Cells were infected with Ot strain UT76 at an MOI of 300:1 (yielding an actual MOI of 3:1 based on image analysis) and incubated at 35 °C for 6 hours. Following infection, cultures were rinsed once, fixed with 4% formaldehyde, and subsequently labeled with Hoechst (nuclei), Evans Blue (cell outlines), and a rat anti-TSA56 monoclonal antibody (Ot). Imaging was performed using a confocal microscope and analyzed with CellProfiler. For Ot entry, we quantified infection by the number of intracellular Ot per infected cell (Bac/Inf). For Ot translocation, we assessed the number of bacteria adjacent to the nucleus per infected cell (NucBac/Inf).

Non-Toxic Drug Doses Selected for Evaluation

| Pathway (S2 Table, sheet “PPI Inhibition”) | Drug | Concentration |
| --- | --- | --- |
| Tricarboxylic acid cycle enzyme complex | AGI-6780 | 16 μM |
| | Devimistat | 20 μM |
| Clathrin-mediated endocytosis, Endocytosis | Dynasore | 32 μM |
| Ubiquitin-dependent protein catabolic process | Bortezomib | 0.01 μM |
| | Carfilzomib | 0.1 μM |
| | MG-132 | 0.2 μM |
| | Ixazomib | 0.04 μM |
| GPCR ligand binding | Maraviroc | 0.1 μM |
| Endoplasmic reticulum to Golgi vesicle-mediated transport | Thapsigargin | 1 nM |
| Microtubule-based process/<br>Microtubule | Vincristine | 0.04 μM |
| | Nocodazole | 1 μg/ml |
| | Colchicine | 1 μg/ml |
| Purine nucleotide biosynthetic process/Purine metabolism | Azathioprine | 12 μM |
| Regulation of actin cytoskeleton/Actin cytoskeleton | CK-636 | 5 μM |
| Arachidonic acid metabolism | Varespladib | 1 μM |
| Cellular response to cytokine stimulus | MRT67307 | 0.2 μM |
| | Ruxolitinib | 10 μM |
| | Tofacitinib | 0.4 μM |

### Secondary siRNA screening

Eleven hits identified from the primary siRNA screen were selected and obtained from Dharmacon. Transfections were performed using 25 nM siRNA and 0.5 µL Lipofectamine 3000 in a total volume of 50 µL Opti-MEM. The transfection mixture was incubated at room temperature for 20 minutes before seeding 8,000 HeLa cells (passage number ≤ 20) in DMEM supplemented with 5% FBS into each well of an 8-well iBidi-treated slide. Cells were incubated at 37 °C in 5% CO_2_ for 24 hours, after which the transfection solution was replaced with fresh DMEM containing 5% FBS. After an additional 24-hour incubation, cells were infected with Ot strain UT76 at an MOI of 1500:1 (yielding an actual MOI of 22:1 based on image analysis). Following a 30-hour infection period, the cultures were rinsed and fixed. Staining was performed as described for the primary screen, and imaging was conducted using a confocal microscope. For Ot entry, we quantified infection by the number of intracellular Ot per infected cell (Bac/Inf). For Ot translocation, we assessed the number of bacteria adjacent to the nucleus per infected cell (NucBac/Inf).

### Statistical tests

All data are expressed as mean ± standard deviation (SD) unless otherwise stated. Statistical analyses were performed using GraphPad Prism software version 11.0.0. Additional data processing and analyses were performed using MATLAB version R2025a (MathWorks) and Python version 3.11. Differences between experimental groups and controls were assessed using unpaired Student’s t-tests, with a p-value < 0.05 considered statistically significant. In drug screening and secondary siRNA screening experiments, fold changes in parameters, including the number of intracellular Ot per infected cell (Bac/Inf) and the number of bacteria adjacent to the nucleus per infected cell (NucBac/Inf), were computed relative to controls, with significance determined by unpaired t-tests. For comparisons of bacterial burden across cell cycle phases in Fig. 5C, statistical significance was assessed using the Mann–Whitney U test, as indicated in the figure legend. These statistical approaches ensured rigorous quantification and validation of our screening data.

## Acknowledgements

We are grateful to lab members at MORU, Cambridge and SiCORE for Systems Pharmacology for their generous support and input. This work was funded by a joint MRC-NSTDA grant awarded to JS and SS, as well as a Royal Society Dorothy Hodgkin Research Fellowship (JS) and Wellcome Trust Senior Research Fellowship (JS).

## Dataset descriptions

S1 Table. Sheet “Screen Results”: Screen results.

S1 Table. Sheet “Toxic Genes”: Toxic genes.

S2 Table. Sheet “GO Inhibit Top100”: Top 100 GO terms for inhibition phenotypes.

S2 Table. Sheet “PPI Inhibit”: PPI networks for inhibition phenotypes.

S2 Table. Sheet “GO Enhance Top100”: Top 100 GO terms for enhancement phenotypes.

S2 Table. Sheet “PPI Enhance”: PPI networks for enhancement phenotypes.

S3 Table. Sheet “Local SSMD Threshold”: Local SSMD thresholds of siRNA plates.

S3 Table. Sheet “Number of Hits”: Number of hits per siRNA plate.

S3 Table. Sheet “Selected Parent Functions”: Selected parent functions.

S3 Table. Sheet “R-square 10-Fold CV”: R² values from 10-fold cross-validation.

---

**Supp. Fig S1:**
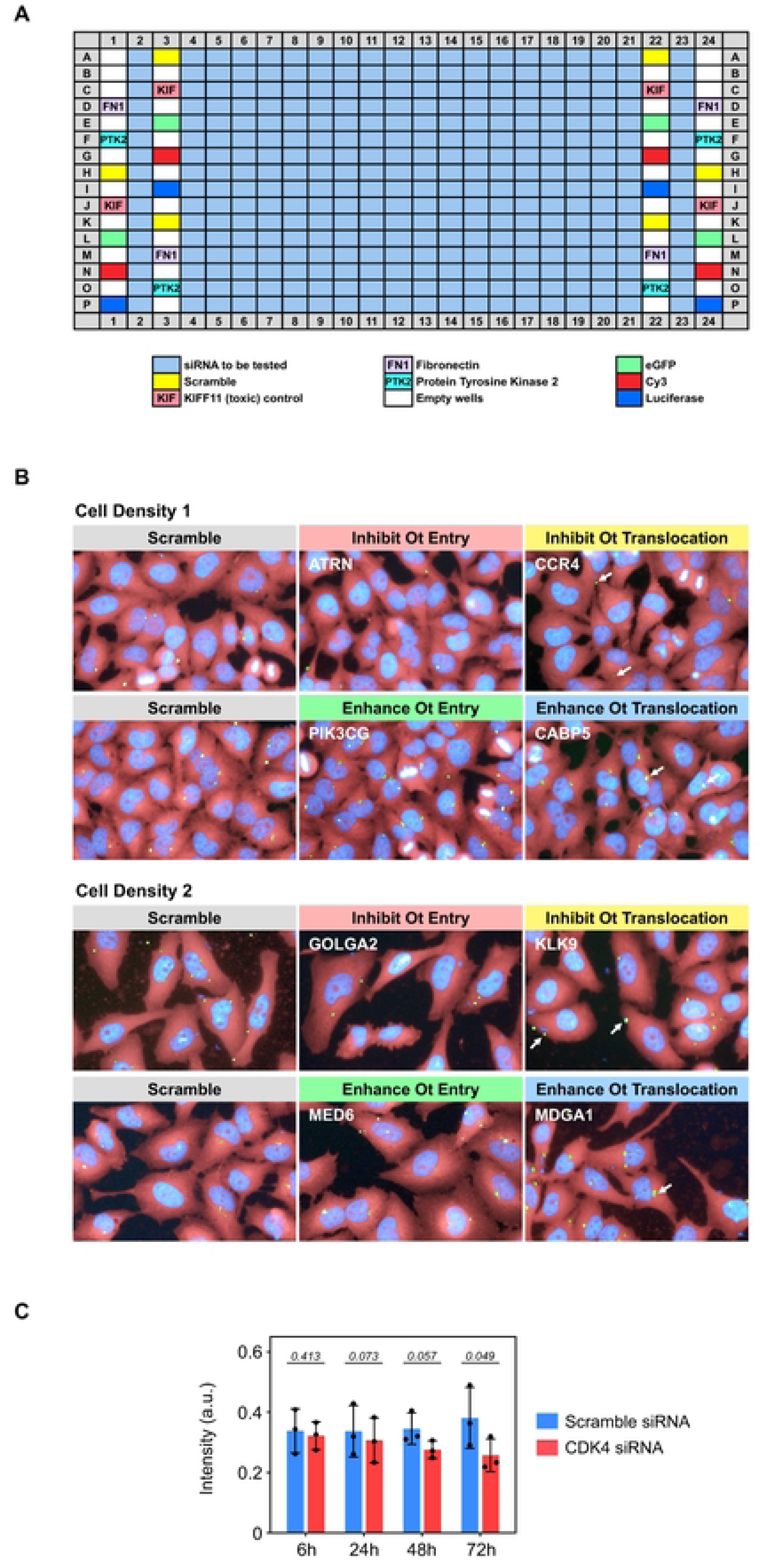
RNAi screen to identify host factors involved in Ot infection. **A.** 384-well plate map showing the layout used for the siRNA screen. **B.** Representative images of screen hits for the four defined phenotypes, with two examples at different cell densities. Images display Hoechst staining (nuclei), FITC labeling (Orientia UT76 protein), and Alexa Fluor 647 labeling (HeLa cells via Evans Blue). **C.** Validation of siRNA-mediated knockdown by immunofluorescence. HeLa cells grown in 8-well ibidi slides were transfected with either scramble control or CDK4 siRNA. Cells were fixed at 6, 24, 48, and 72 hours post-transfection, labeled with anti-CDK4 antibodies, and imaged by confocal microscopy. Fluorescence intensity was quantified using CellProfiler, and statistical significance was determined by Student’s t-test (GraphPad Prism).

**Supp. Fig. S2.**
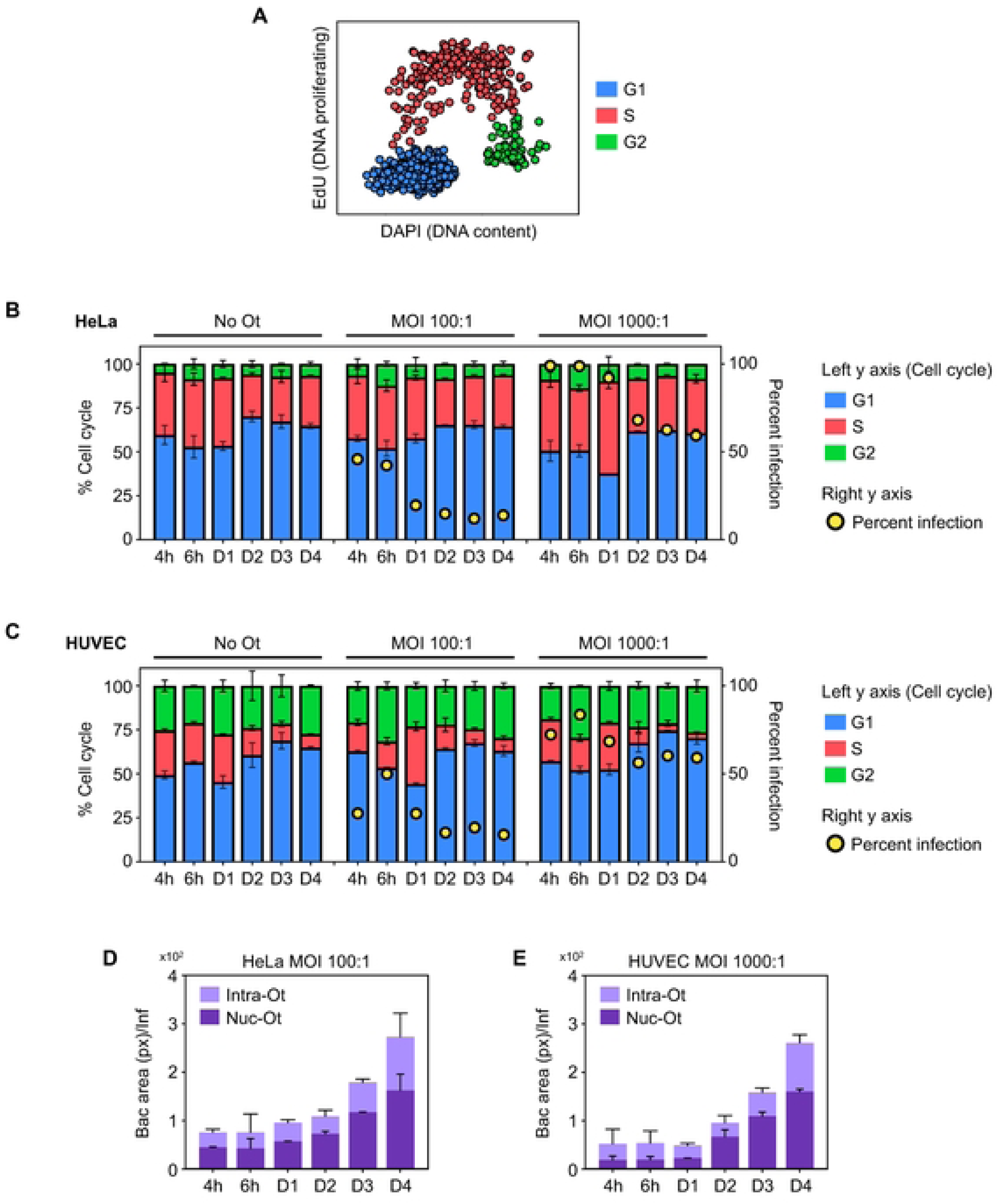
Host Cell Cycle Analysis and Ot Colony Dynamics. **A.** Cell cycle plot of HeLa cells stained with EdU (marking replicating DNA) and Hoechst (staining total DNA), enabling classification into G1 (blue), S (red), and G2 (green) phases. Data represent single-cell-level measurements. **B-C.** Quantification of cell cycle distribution into G1 (blue), S (red), and G2 (green) phases and percentage of Ot-infected cells (yellow dot) in HeLa (**B**) and HUVEC (**C**) over time (4-6 hours, 1, 2, 3, and 4 days post-infection). **D-E.** Quantification of intracellular Ot (Intra-Ot) and perinuclear Ot (Nuc-Ot) area retrieved from infected cells of HeLa (MOI 100:1, **D**) and HUVEC (MOI 1000:1, **E**) cells at the corresponding time points.

